# Joint Distribution of Protein Concentration and Cell Volume Coupled by Feedback in Dilution

**DOI:** 10.1101/2023.04.27.538534

**Authors:** Iryna Zabaikina, Pavol Bokes, Abhyudai Singh

## Abstract

We consider a protein that negatively regulates the rate with which a cell grows. Since less growth means less protein dilution, this mechanism forms a positive feedback loop on the protein concentration. We couple the feedback model with a simple description of the cell cycle, in which a division event is triggered when the cell volume reaches a critical threshold. Following the division we either track only one of the daughter cells (single cell framework) or both cells (population framework). For both frameworks, we find an exact time-independent distribution of protein concentration and cell volume. We explore the consequences of dilution feedback on ergodicity, population growth rate, and the bias of the population distribution towards faster growing cells with less protein.

## 1 Introduction

Due to low copy number of molecules, gene expression is a stochastic process [1,21]. The typical question asked of a mathematical model is to determine the distribution of a protein level at steady state [3,45]. Traditionally, the number of cells is treated as a discrete variable which is subject to production and depletion events [4,48]. Protein production is modelled by a compound Poisson process [12,34]. Production events are referred to as bursts and the number of protein molecules produced per event as the burst size [6,7]; these are assumed to be independent and identically geometrically distributed [35,36]. Depletion of protein can be driven by two different mechanisms: degradation or partitioning at cell division [30].

Degradation removes one molecule at a time and leads to tractable models with explicit steady state distributions [13,24,32]. Partitioning based depletion requires that a model for the cell cycle be combined with the model for gene expression [17,42]. At the end of the cell cycle, each molecule “flips a coin” to decide which of the two daughter cells it goes to [5]. From a single cell line perspective, the partitioning of molecules manifests as a downward jump from the current protein cell count to roughly one half of it. Other types of partitioning than symmetric binomial have also been considered [30]. Models with partitioning errors are more complex than the degradation based models [29,49].

One way to circumvent the complexity due to partitioning is to track the protein concentration instead of the protein count [8,31]. Production is still modelled by a compound Poisson process [28,33], but the geometric distribution of burst sizes is replaced by its coarse-grained continuous counterpart, the exponential distribution [11]. Following other authors, we thereby assume that burst sizes are in units of concentration [22]; this implies a coupling of burst size to cell volume as shown experimentally [40]. The binomial partitioning reduces, by the law of large numbers, to the exact halving of molecules [27]. At the same time, the volume of the mother cell is assumed to split into two exact halves [46,53,54]. Consequently, the concentration (the number of proteins per unit volume) is unaffected by the partitioning at the end of the cell cycle. Instead, it decays continuously over the duration of the cell cycle as the volume of the cell grows [15,52]. In the simplest case, we assume that the volume grows exponentially during the cell cycle, so that the concentration decays exponentially. The basic model without feedback leads to the gamma distribution of protein concentration [10,25,39].

We consider a scenario where the level of a given protein negatively regulates the exponential rate of cell volume expansion [47,55]. Such feedback coupling is often manifested as a result of the burden placed by the protein on the cells’ expression/metabolic machinery [16,37,41,51]. In the feedback scenario, it is important to distinguish between the single cell (genealogy) framework and the population framework [19,23,50]. For a single cell line case, the model is a bivariate Markov process, the variables being the protein concentration and cell volume. For the population case, the model is a measure-valued Markov process [14], in which the state of the population is represented by a measure (a generalised density) on the two-dimensional space of possible protein concentrations and cell volumes. Section 2 introduces the two frameworks in detail and states the main results. The results are derived in Sections 3 and 4 by studying the associated evolution equations (Chapman–Kolmogorov and the population balance equations). Section 5 highlights the implications of the analysis.

## 2 Model Formulation

For each cell, we track its concentration *x*(*t*) and volume *v*(*t*) at time *t*. Protein synthesis occurs in random discrete events with stochastic rate *α*. Therefore, the interarrival times of synthesis events are i.i.d. exponentially distributed random variables with mean 1*/α*. Each burst event creates a jump or burst in the protein concentration *x* → *x* + *B*, where the burst size *B* is drawn from exponential distribution with mean *β*. The cell volume is not affected by a protein synthesis burst.

Between successive bursts, the cell volume increases with a concentration dependent rate and the concentration is diluted according to differential equations

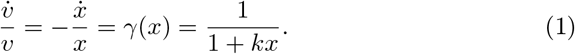

The volume *v*(*t*) is strictly increasing during the cell cycle. The concentration *x*(*t*) is strictly decreasing between consecutive production bursts. When the cell volume reaches a given threshold 2*v*^∗^, a cell division event occurs, and the volume is immediately halved

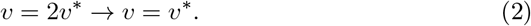

Cell division does not affect the protein concentration. Typical protein and volume trajectories of a single cell line are shown in Figure 1. The reset rule (2) is based on cell division as a Sizer [20].

**Fig. 1.**
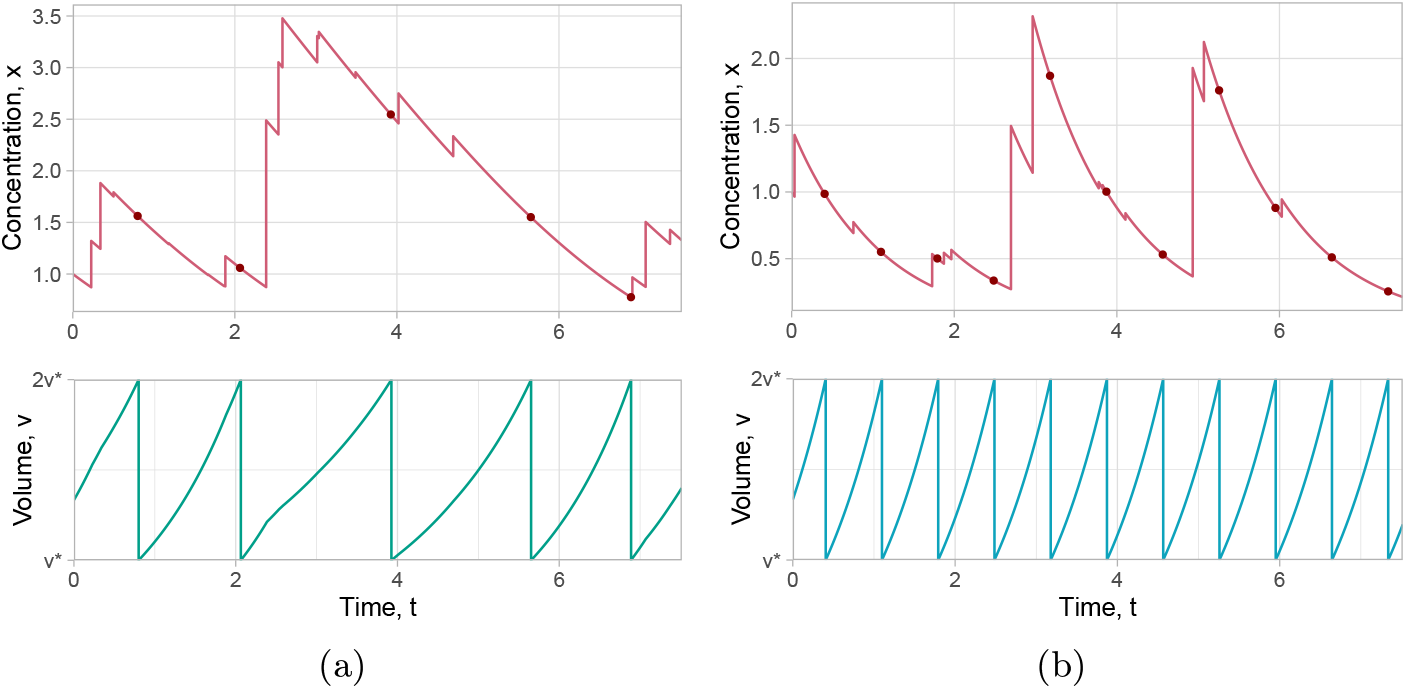
Sample trajectories of the concentration *x*(*t*) and the volume *v*(*t*) in a single cell affected by (a) the strong feedback with *k* = 2, (b) the absence of the feedback, i.e., *k* = 0; other parameters of simulations are *α* = 0.9, *β* = 5. Each division event is marked by a red dot on the concentration trajectory, which corresponds to an instant drop from 2*v*^∗^ to *v*^∗^ on the volume trajectory.

### 2.1 Single Cell Model: a Bivariate Markov Process

In the single-cell model, we are looking at a single cell line: we do not follow the other daughter cell that is created in a cell division. The model is then a piecewise deterministic bivariate Markov process 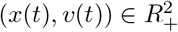. Let *f* (*x, v, t*) denote the probability density function (pdf) of the random vector (*x*(*t*), *v*(*t*)) at time *t*. We have the initial condition *f* (*x, v, t* = 0) = *δ*(*x* − *x*_0_)*δ*(*v* − *v*_0_), where *x*_0_ *>* 0 and *v*_0_ *>* 0 are known initial concentration and volume. We thereby assume that the initial volume is less that the critical volume at which a cell division event is triggered: *v*_0_ *<* 2*v*^∗^.

For *t >* 0, the pdf *f* (*x, v, t*) satisfies a bivariate Chapman–Kolmogorov equation, which is formulated in Section 3. We solve the Chapman–Kolmogorov equation at steady state. We find that the stationary distribution has the product form *f* (*x, v*) = *p*(*x*)*g*(*v*), where the marginal distributions are given by

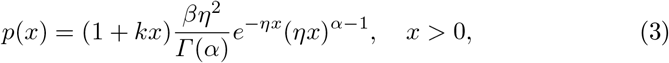

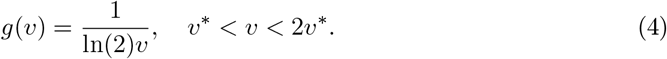

The protein distribution (3) exists only if *η* = 1*/β* − *αk >* 0.

In the absence of feedback (*k* = 0), the cell cycle length is equal to the doubling time *T* = ln 2 of the exponential function, and the cell volume *v*(*t*) is a *T* -periodic function (Figure 1b). Because of the periodicity, the time-dependent pdf *f* (*x, v, t*) does not converge to the stationary distribution *p*(*x*)*g*(*v*) if *k* = 0. The inclusion of feedback breaks the periodicity (Figure 1a). Therefore, for *k >* 0,

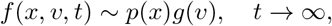

i.e. the time-dependent distribution converges to the stationary distribution in the large-time limit (the ergodic property).

### 2.2 Population Model: a Measure-valued Markov Process

In the population model, the end of the cell cycle triggers a branching (division) event: the current process (the mother cell) is terminated and replaced with two new processes (daughter cells), which inherit the mother’s protein concentration, and half of the mother’s cell volume. The daughter processes evolve henceforth independently of each other (Figure 2, left panel).

**Fig. 2.**
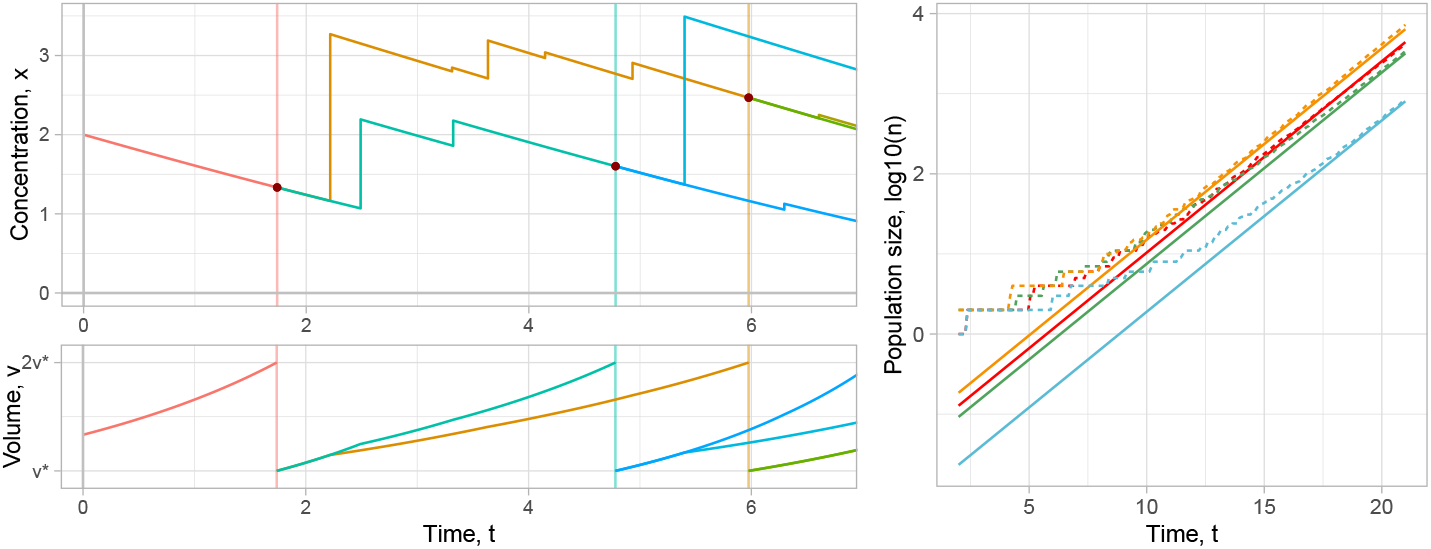
*Left*: the time evolution of a sample cell population, where an individual color is assigned to each cell. We plot the protein concentration in the cell, its volume, and the division time (vertical lines). *Right:* The time evolution of the log-scaled sample population size in four different simulation runs with the same initial conditions and parameters values *α* = 0.9, *β* = 0.5, *k* = 2, and *v*^∗^ = 1.5 (dashed lines); the solid lines correspond to the exponential growth with the rate constant (8), shifted by the random factor log_10_ *W* (*x*_0_, *v*_0_), whose specific values were estimated for each simulation using linear regression. Each simulation has random fluctuations at the beginning and afterwards converges to the expected exponential growth.

The above recursive construction leads to a sequence of bivariate Markov processes

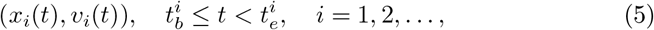

where 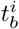 and 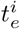 denote the beginning and the end of the cell cycle of the *i*th cell, and *x*_*i*_(*t*) gives the protein concentration at time *t* of the *i*th cell and *v*_*i*_(*t*) gives its volume. The original cell from which the entire population is derived is indexed by *i* = 1. The ordering of the rest of the sequence depends on the algorithmic implementation of the population process and is immaterial for our purposes.

The composition of the population at time *t* can be represented by the empirical population density (or measure)

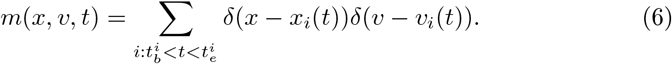

The sum in (6) extends over the cells that exist at time *t*. Each existing cell contributes to the empirical population density (6) by a unit mass placed on top of (*x*_*i*_(*t*), *v*_*i*_(*t*)) ∈ (0, ∞) × [*v*^∗^, 2*v*^∗^]. The integral of the empirical population density over the whole state space,

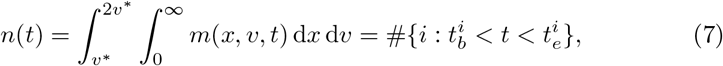

gives the total number of cells existing at time *t* and is finite. The empirical population density (6) is an example of a measure-valued Markov process [14,18].

Let us consider the expected value of the empirical population density

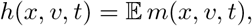

the expectation being taken over all possible realisations of the population process. At the initial time *t* = 0, we have a single cell with non-random initial concentration *x*_0_ and initial volume *v*_0_, so that

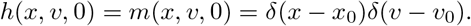

For *t >* 0, the expected population density *h*(*x, v, t*) satisfies a population balance equation. This is formulated in Section 4.

The large-time behaviour of the population balance equation is characterised by its principal eigenvalue *λ*, the associated eigenfunction *f* (*x, v*), and the adjoint eigenfunction *w*(*x, v*). In Section 4, we show that

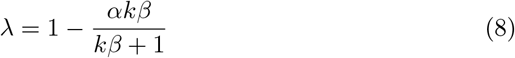

and *f* (*x, v*) = *p*(*x*)*g*(*v*), where

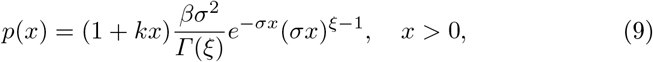

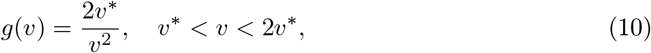

where *ξ* = *α/*(1+*kβ*) and *σ* = 1*/β* − *αk/*(1+*kβ*). The principal eigenfunction (9) exists (in the sense of belonging to the space of integrable functions) only if *σ >* 0. We also see that *σ >* 0 is equivalent to *λ >* 0. The condition *σ >* 0 is weaker than the condition *η >* 0 for the existence of the stationary distribution (3).

Spectral decomposition implies that the expected (non-random) population density satisfies

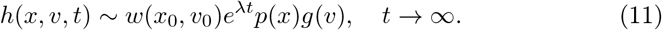

The adjoint eigenfunction *w*(*x*_0_, *v*_0_) *>* 0, whose functional form we do not determine, thus characterises the influence of the initial condition on the large-time asymptotics (11). Analogously to what was said in the single cell model, (11) holds only in the aperiodic case (*k >* 0).

By the theory of supercritical branching processes [2,26], the (random) empirical population density satisfies

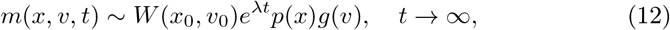

where *W* (*x*_0_, *v*_0_) *>* 0 is a random variable dependent on initial data such that 𝔼 *W* (*x*_0_, *v*_0_) = *w*(*x*_0_, *v*_0_). Equation (12) can equivalently be written in terms of the population size (7) and the normalised protein distribution as

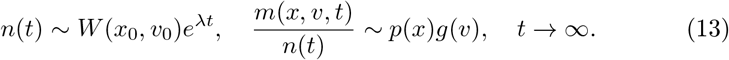

By the first relation in (13), the population *n*(*t*) increases eventually exponentially with the rate constant (8); the initial condition and the low-population noise affects the large-time behaviour of the population only through the random pre-exponential factor *W* (*x*_0_, *v*_0_) (Figure 2, right panel). By the second relation in (13), the distribution of protein concentration and cell volume among a large population is (nearly) nonrandom and given by (9)–(10) (Figure 3).

**Fig. 3.**
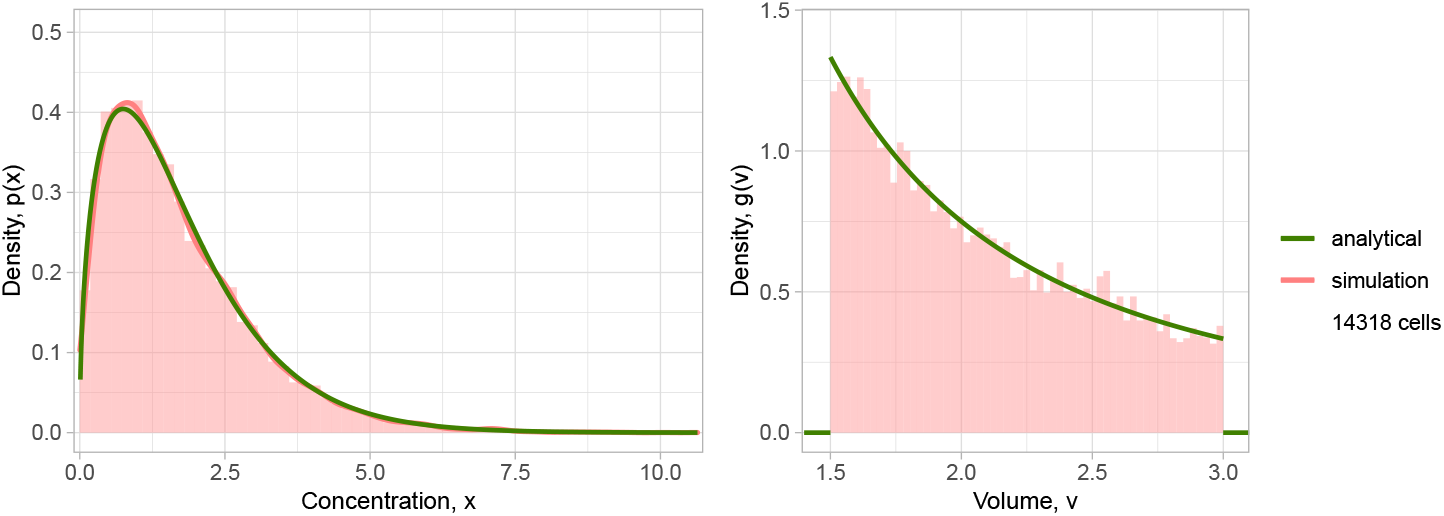
Comparison of the analytical distributions (9) and (10) to the results of a large-time kinetic Monte Carlo simulation with parameters *α* = 2, *β* = 0.45, *k* = 0.7, *v*^∗^ = 1.5, a simulation endpoint *t* = 20, and initial conditions *x*_0_ = *v*_0_ = 2.

## 3 Chapman–Kolmogorov Equation

The goal of this section is to derive the single-cell stationary distributions *p*(*x*) of protein concentration *x* and *g*(*v*) of cell volume *v*. We start by formulating the time-dependent problem. The probability for the cell to be of volume *v*^∗^ ≤ *v* ≤ 2*v*^∗^ and to have the protein concentration *x >* 0 at time *t >* 0 is given by the joint probability density function is *f* (*x, v, t*); its time evolution is described by Chapman-Kolmogorov equation:

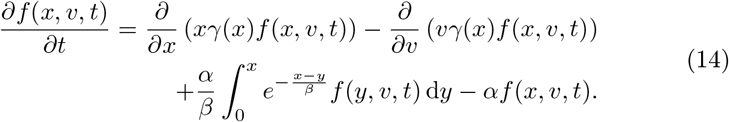

The initial and boundary conditions are:

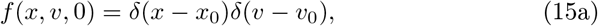

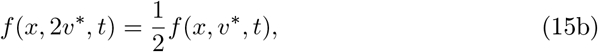

where *x*_0_ is initial protein concentration in the cell, *v*_0_ is its initial volume. The Chapman–Kolmogorov equation (14) is a partial integro-differential equation. The differential operator on the right hand side of (14) drives the drift of the probability mass due to the deterministic flow (1). The integral operator in (14) provides the transfer of probability mass due to instantaneous protein bursts. The boundary condition (15b) captures the halving of cell volume in cell division. Integrating (14) over the state space (*x, v*) ∈ (0, ∞) × [*v*^∗^, 2*v*^∗^] confirms that the total probability 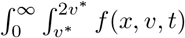 d*x* d*v* remains constant over time for solutions *f* (*x, v, t*) to (14). The boundary fluxes thereby cancel thanks to the boundary condition (15b).

Since our aim is to find a stationary distribution *f* (*x, v*), we set ∂*f/*∂*t* = 0 and subsequently apply the Leibniz integral rule to the last two terms, which yields an integral equation:

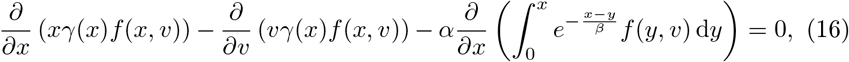

the boundary condition for which is the same as (15b).

We use the Fourier method of separation variables, i.e. we assume that *f* (*x, v*) can be represented as a separable function

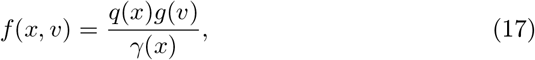

where *γ*(*x*) is given by (1) and *q*(*x*) and *g*(*v*) are to be determined. The marginal protein stationary distribution is then *p*(*x*) = *q*(*x*)*/γ*(*x*) and that of the cell volume is *g*(*x*). We substitute (17) into (16) and rearrange the result so that on the left-hand side all terms depend only on the variable *x*, and on the right-hand side only on *v*:

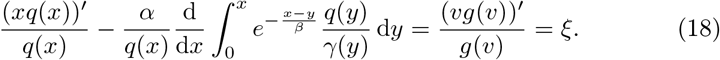

Since both sides are independent from one another, each side must be equal to a constant, which we denoted as *ξ*. We rewrite (18) and add the boundary condition on *g*(*v*):

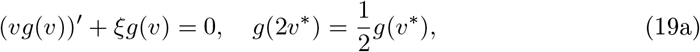

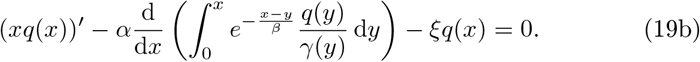

First, we solve equation (19a), which is an ordinary differential equation of the first order. The general solution is

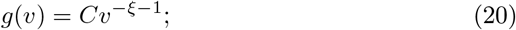

upon applying the boundary condition, we find that *ξ* = 0. Since *g*(*v*) is a probability distribution, the integral of *g*(*v*) over the interval (*v*^∗^, 2*v*^∗^) must be equal to one. The stationary distribution of the cell volume is thus (4).

We proceed with the second equation (19b); after we substitute *ξ* = 0, it reduces to :

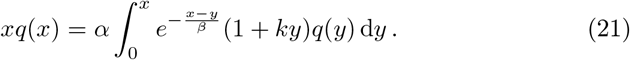

The homogeneous Volterra integral equation (21) be solved by standard methods [55]. Solving and returning to the protein distribution via *p*(*x*) = *q*(*x*)*/γ*(*x*) leads to (3).

## 4 Population Balance Equation

The goal of this section is to derive the whole-population time-invariant distributions *p*(*x*) of protein concentration *x* and *g*(*v*) of cell volume *v* as well as the large-time population growth rate constant *λ*. We start by formulating the time-dependent problem. The expected population density *h*(*x, v, t*) of cells with concentration *x >* 0 and volume *v*^∗^ ≤ *v* ≤ 2*v*^∗^ at a particular point of time *t* ≥ 0 satisfies a population balance equation. In the interior of the state space (*x >* 0, *v*^∗^ *< v <* 2*v*^∗^), the population balance equation coincides with the Chapman–Kolmogorov equation (14):

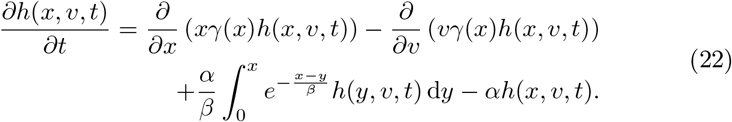

The initial and boundary conditions are:

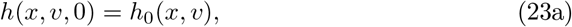

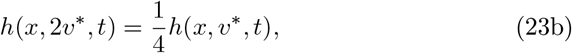

where *h*_0_(*x, v*) in (23a) is an initial population density. The extra factor of two in the boundary condition (23b) compared to the previous boundary condition (15b) reflects the cell doubling at the end of a cell cycle in the population scenario.

The population grows in time, and we assume that *h*(*x, v, t*) has the following separable form:

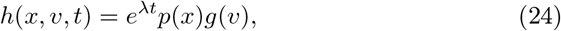

where

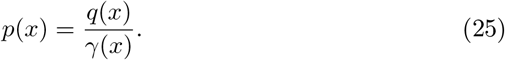

The function *γ*(*x*) is given by (1) while *λ, q*(*x*), and *g*(*v*) are to be determined. Substituting (24) and (25) into (22) and (23b) yields

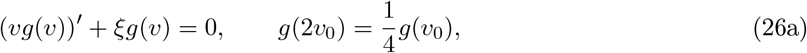

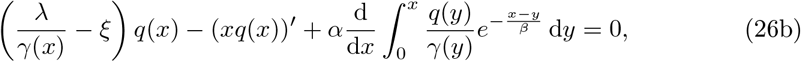

where *ξ* is a constant independent of *x* and *v*. Equation (26a) is equivalent to (19a) except the boundary condition. The general solution remains to be (20), but the boundary condition now yields *ξ* = − 1; from this, the population volume distribution (10) follows immediately.

Substituting *ξ* = −1 into (26b), we obtain the integro-differential equation

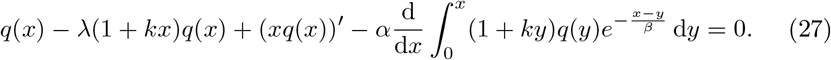

Applying the Laplace transform to (27) yields

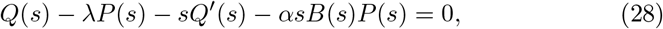

where *P* (*s*) and *Q*(*s*) are the Laplace images of functions *p*(*x*) and *q*(*x*), respectively, and 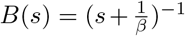 is the Laplace transform of the complementary cumulative distribution function of the (exponential) burst size distribution. Applying the Laplace transform directly to (25), one obtains

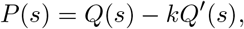

which is used to rewrite (28) into an ODE for *Q*(*s*):

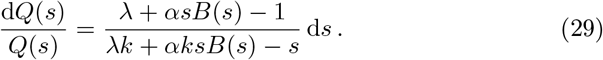

In the case of exponentially distributed burst sizes, the right-hand side of (29) simplifies to a rational fraction:

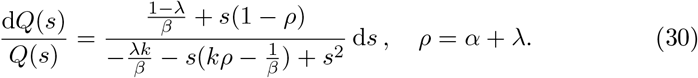

The solution approach requires the partial fraction decomposition of the right-hand side of (30). The quadratic in the denominator has two real roots

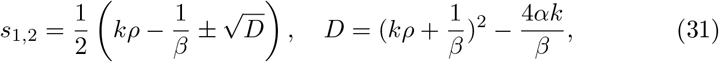

where *s*_1_ is strictly positive and *s*_2_ is strictly negative for any positive values of parameters *α, β, λ* and *k*. The decomposition of (30) takes the form:

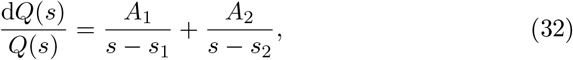

where *A*_1_ and *A*_2_ are defined by

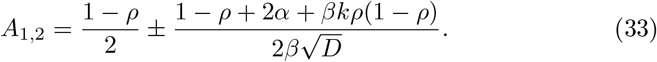

The solution of the ODE (32) is

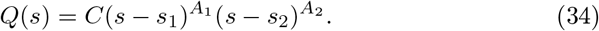

The Laplace transform (34) has to be analytic in the complex half-plane, i.e., Re(*s*) *>* 0. Therefore, *A*_1_ ∈ *{*0, 1, 2, *}*. In particular, the principal eigenvalue is obtained by setting

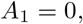

which implies

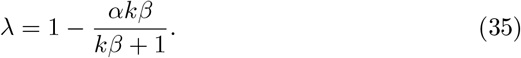

We expect that setting *A*_1_ = 1, 2, … leads to additional eigenvalues.

We substitute (35) into (31) and (33) and simplify; the resulting values are strictly negative, and we introduce additional symbols for their opposite values, which are strictly positive:

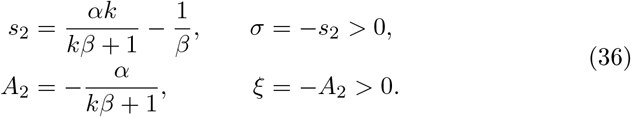

Inserting *s* = 0 into (28) and using the normalisation condition *P* (0) = 1 yield

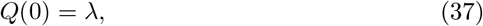

which is used to find the value of *C* in (34). Applying the inverse Laplace transform to (34), we obtain

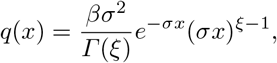

from which, by relation (25), the population protein concentration distribution (9) immediately follows.

## 5 Conclusions

In this paper, we studied a stochastic model for a protein that inhibits the growth of cell volume. The main result is the large-time distribution of protein concentration and cell volume in a single cell over time (3)–(4) and in an expanding cell population (9)–(10). Interestingly, the two are independent in the large-time limit (but interdependent transiently in the presence of the dilution feedback). We expect that the large-time independence carries over to more complex models of cell division than the reset rule (2). However, additional coupling between protein and cell size (e.g. volume-dependent production, partitioning noise) may introduce a dependence between the two variables [38].

The single-cell stationary distribution (3)–(4) exists if the product *αβ* of burst frequency and burst size is less than the maximal dilution rate 1*/k* = lim_*x*→∞_ *x/*(1 + *kx*). Clearly, in the alternative case (*αβ >* 1*/k*), the build up of protein prevents stationarity [55]. In the population scenario, the large-time distribution (9)–(10) exists if the large-time population growth rate constant *λ* (8) is positive. In the alternative scenario ((*α* − 1)*β >* 1*/k*), the build up of protein overburdens the cells and stalls the population growth. We note that this cannot happen in the low burst frequency (high noise) scenario *α <* 1.

In the absence of dilution feedback, the volume process is periodic and hence does not converge to its stationary distribution (Figure 1b). Periodicity also appears for more complex cell division mechanisms than (2) as long as (i) the volume grows exponentially and (ii) the cell divides into two equal halves [9]. Feedback in dilution makes the growth protein-dependent and hence non-exponential. As a consequence, we get rid of the periodicity (Figures 1a and 2) and obtain ergodicity i.e. convergence of the large-time distribution (Figure 3).

Figure 4 visualises the effects of dilution feedback and population framework on the protein and volume distribution. Inclusion of feedback tilts the concentration distribution to the right (Figures 4a and 4c). Inclusion of feedback does not affect the volume distribution (Figures 4b and 4d). Population volume distribution is tilted to the left compared with the single cell distribution (Figures 4b and 4d). Without feedback, the concentration distribution is the same gamma distribution in a population and for a single cell (Figure 4a). With feedback, the population distribution is tilted to the left compared to the single cell distribution (Figure 4c). Consequently, the fraction of cells above a concentration threshold in the population is smaller than the fraction of time a single cell has concentration above the threshold. This has important consequences for drug-tolerant persisters in microbial and cancer cells that will be rarer than as predicted by classical simulation if the feedback is present [43,44].

**Fig. 4.**
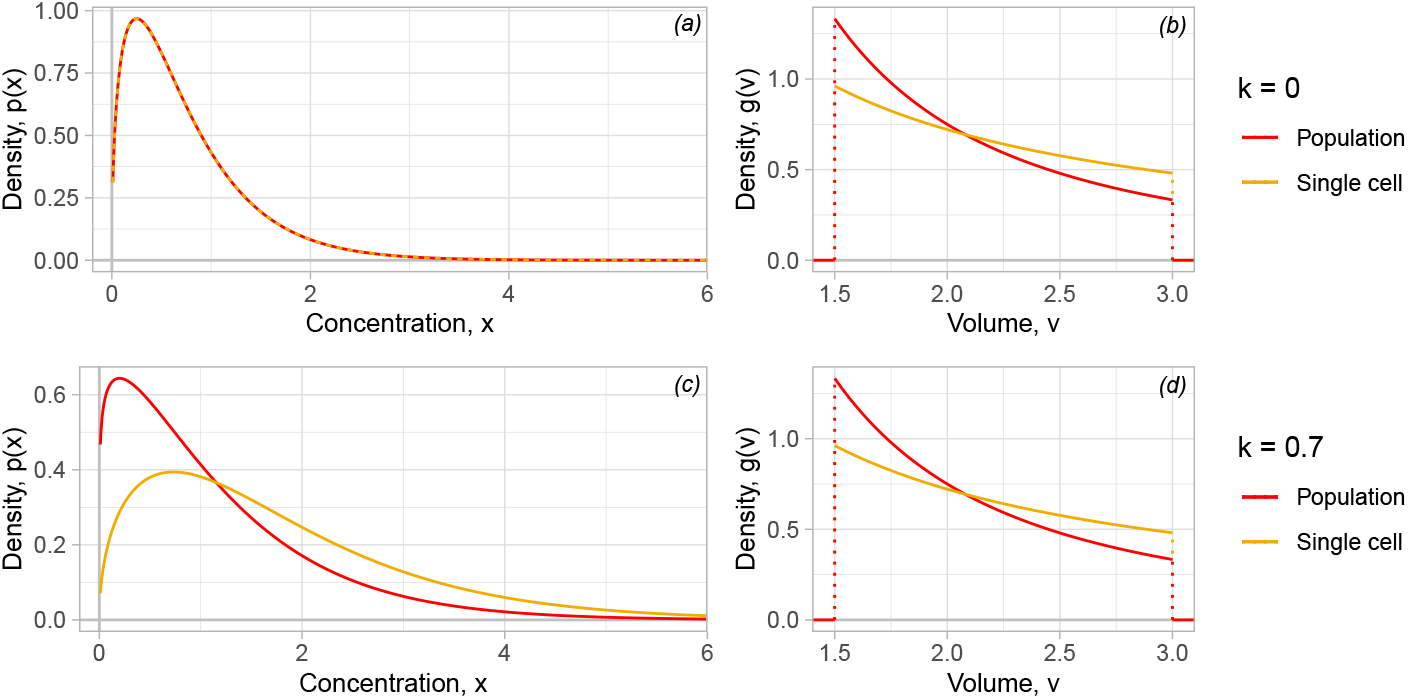
The effect of absence (the first row) and presence (the second row) of feedback in dilution on the large-time distributions of protein concentration *x* and cell volume *v*. Parameters are as follow: *α* = 1.5, *β* = 0.5, *k* = 0.6, *v*^∗^ = 1.5.

## Acknowledgements

PB was supported by the Slovak Research and Development Agency under contract no. APVV-18-0308, and VEGA grants 1/0339/21 and 1/0755/22.

